# Integrating gene expression with summary association statistics to identify susceptibility genes for 30 complex traits

**DOI:** 10.1101/072967

**Authors:** Nicholas Mancuso, Huwenbo Shi, Pagé Goddard, Gleb Kichaev, Alexander Gusev, Bogdan Pasaniuc

## Abstract

Although genome-wide association studies (GWASs) have identified thousands of risk loci for many complex traits and diseases, the causal variants and genes at these loci remain largely unknown. We leverage recently introduced methods to integrate gene expression measurements from 45 expression panels with summary GWAS data to perform 30 transcriptome-wide association studies (TWASs). We identify 1,196 susceptibility genes whose expression is associated with these traits; of these, 168 reside more than 0.5Mb away from any previously reported GWAS significant variant, thus providing new risk loci. Second, we find 43 pairs of traits with significant genetic correlation at the level of predicted expression; of these, 8 are not found through genetic correlation at the SNP level. Third, we use bi-directional regression to find evidence for BMI causally influencing triglyceride levels, and triglyceride levels causally influencing LDL. Taken together, our results provide insights into the role of expression to susceptibility of complex traits and diseases.

## Introduction

Although genome-wide association studies (GWASs) have identified tens of thousands of common genetic variants associated with many complex traits^1^, with some notable exceptions^2; 3^, the causal variants and genes at these loci remain unknown. Multiple lines of evidence show that GWAS risk variants co-localize with genetic variants that regulate expression—i.e. expression quantitative trait loci (eQTL)^4^. This suggests that a substantial proportion of GWAS risk variants influence complex trait by regulating gene expression levels of their target genes^4-7^. Analyses of genotype, phenotype, and gene expression measurements from multiple tissues in the same set of individuals can directly investigate this plausible chain of causality. However, doing so is challenging due to cost and tissue availability; therefore, GWAS and eQTL data sets remain largely independent (i.e. no overlapping subjects)^8; 9^. Recent work demonstrated that using eQTL data to predict expression into the much larger GWAS followed by association testing can identify new susceptibility genes^10-12^. This approach, referred to as transcriptome-wide association study (TWAS), provides testable hypotheses under the molecular cascade of genetic variation impacting expression which in turn impacts complex trait.

In this work we connect TWAS to a test for non-zero genetic covariance between expression and trait, and extend it to estimate the genetic correlation between expression and trait. This interpretation enables us to develop new methods that characterize the relationship between complex traits using gene effects instead of single nucleotide polymorphism (SNP) effects. In particular, we estimate the genetic correlation between pairs of traits at the level of predicted expression; this is analogous to computing genome-wide genetic correlation between traits^13^, with correlations being determined over gene effects rather than SNP effects. Finally, we use a bi-directional regression approach^14^ to investigate putative causal direction for pairs of traits. This approach compares models that regress over estimated effects for identified susceptibility genes and is conceptually similar to recent work^15^ which uses effects of GWAS risk SNPs.

We analyze 30 GWASs spanning over 2.3 million phenotype measurements^16-29^ jointly with 45 expression panels sampled from more than 35 tissues and perform 30 TWASs to gain insights into the role of expression in complex trait etiology. First, we identify 1,196 genes associated with these complex traits and diseases resulting in 1,789 distinct gene-trait pairs. Of these pairs, 168 did not overlap (0.5Mb from TSS) a genome-wide significant SNP for that respective trait, which we consider to be novel risk loci. We also find 219 cases where association signal is stronger in TWAS suggesting that allelic heterogeneity plays a role in regulating expression. Consistent with previous reports^11; 12^, the vast majority of susceptibility genes were not proximal to the GWAS index SNP. Second, we estimate genetic correlation between these traits at the level of predicted expression and identify 43 pairs with significantly non-zero estimates; of these, 35 can be identified through genetic correlation analyses at the SNP level with 8 being identified only by analyzing predicted expression. These results suggest that a significant component of genetic correlation between complex traits can be explained by predicted expression. Lastly, we perform bi-directional analyses to provide evidence for putative causal effects between pairs of traits. Using this approach, we find evidence consistent with a causal model where body mass index (BMI) influences triglyceride levels, in line with earlier work^15^. We also report a novel result suggesting that triglyceride levels influence low-density lipoprotein (LDL) levels. Overall, our results shed light on shared biological mechanisms responsible for susceptibility to disease and complex trait, as well as potential downstream effects between traits.

## Methods

### Data Sets

We used summary association statistics from 30 large-scale (N>20,000 subjects) GWAS including various anthropometric^16; 28; 29^ (BMI, femoral neck bone mineral density (BMD), forearm BMD, height, lumbar spine), hematopoietic^24; 27^ (hemoglobin, HBA1C, mean cell hemoglobin (MCH), MCH concentration, mean cell volume, number of platelets, packed cell volume, red blood cell count), immune-related^18; 20^ (Crohn’s disease, inflammatory bowel disease, rheumatoid arthritis, and ulcerative colitis), metabolic^17; 23^ (age of menarche, fasting glucose, fasting insulin, high-density lipoprotein, HOMA-B, HOMA-IR, low-density lipoprotein, triglycerides, type 2 diabetes, and total cholesterol levels), neurological^19^ (schizophrenia), and social phenotypes^22^ (college and educational attainment; see Supplementary Table 1). We removed SNPs that were strand-ambiguous or those with minor allele frequency £ 1% (see Supplementary Table 1).

Gene expression data from RNA-Seq data were obtained from the CommonMind Consortium^30^ (CMC; brain; N=613), the Genotype-Tissue Expression project^8^ (GTEx; 41 tissues; see Supplementary Table 2 for sample size per tissue), Metabolic Syndrome in Men (METSIM; adipose; N=563)^31; 32^. Expression microarray data were obtained from the Netherlands Twins Registry^33^ (NTR; blood; N=1,247), and the Young Finns Study^34; 35^ (YFS; blood; N=1,264).

### Performing TWAS using GWAS summary statistics

We estimated SNP heritability for observed expression levels partitioned into cis-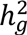 (1 Mb region surrounding the gene) and trans-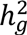 (rest of genome) components. We used the AI-REML algorithm implemented in GCTA^36^, which allows estimates to fall outside of the (0, 1) boundaries to maintain unbiasedness. To control for confounding, we included batch variables and the top 20 principal components estimated from genome-wide SNPs. Genes with significant cis-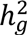 (p < 0.05 in a likelihood ratio test between the cis-only and joint model) in expression data were used for prediction. We performed a prediction-based transcriptome wide association study (TWAS) for each of the 30 GWAS using the summary approach described in ref^11^. In brief, we estimated the strength of association between predicted expression of gene and complex trait (*z_TWAS_*), as function of the vector of GWAS association summary Z-scores at a given cis locus ***z***_*T*_ and the LD-adjusted weights vector learned from the gene expression data ***w***_*GE*_ as

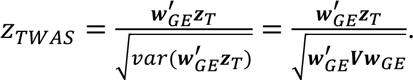

where ***V*** is a covariance matrix across SNPs at the locus (i.e. LD). We estimated ***w***_*GE*_ using GBLUP^37^ from eQTL data and computed *z_TWAS_* using GWAS summary data for all 30 traits and the ∼36k gene expression measurements across all studies. We removed all loci in the human leukocyte antigen (HLA) region due to complex LD patterns. We conservatively account for multiple tests using trait-specific Bonferroni correction factors (see Supplementary Table 2).

### Estimating the proportion of trait variance explained by predicted expression

We use an LD-Score^38; 39^ approach^11^ to quantify the heritability for a complex trait explained by predicted expression (denoted here as 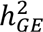). The expected *χ*^2^ statistic under a polygenic trait is 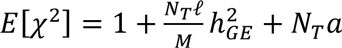 where *N_T_* is the number of individuals in the GWAS, *M* is the number of genes, *ℓ* is the LD-score, and *a* is the effect of population structure. We estimate *ℓ* for each gene by predicting expression for 503 European samples in 1000Genomes^40^ using the BLUP weights (see above) and then computing sample correlation. For each complex trait we perform LD-Score regression using 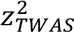 (which is asymptotically equivalent to *χ*^2^) to infer 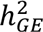. We estimate heritability for each expression study separately, to account for varying sample sizes and repeated gene measurements.

### Estimating genetic correlation of expression and complex trait from summary data

Let expression and trait be modeled as a linear function of the genotypes in a ∼1Mb local region flanking the gene: ***y***_*GE*_ + ***Xβ***_*GE*_ + *∊_GE_* and ***y***_*T*_ + ***Xβ***_*T*_ + *∊_T_* where ***X*** is the standardized genotype matrix, *β_GE_*(*β_T_*) are the standardized effects, and *∊_GE_*(*∊_T_*) is environmental noise for expression (trait). The local covariance between expression and complex trait is

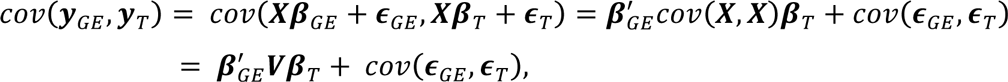

where *V* is the LD matrix. If no individuals are shared between studies then *cov*(*∊_GE_*, *∊_T_*) = 0, (as in eQTL and GWAS studies). The local genetic correlation can be computed as

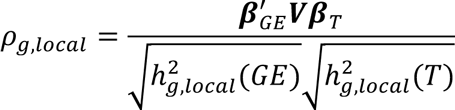

where 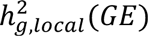 is the local SNP-heritability^41^ for expression (trait) estimated at the locus captured by ***X***; however, this requires knowing the true effect sizes. Previous work^41^ describes a method to obtain unbiased estimates for ***β***_*i*_ using genome-wide association summary statistics (i.e. Z-scores) and reference LD. Given association statistics ***z***_*T*_, an LD-adjusted effect size estimate is computed as 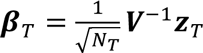. Hence, an estimate of the local genetic covariance^42^ is given by

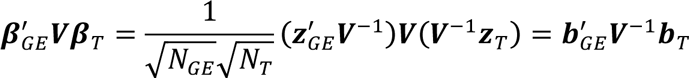

where ***b***_*GE*_(***b***_*T*_) are the marginal (i.e. LD-unadjusted) effect sizes^41; 43^. It follows that

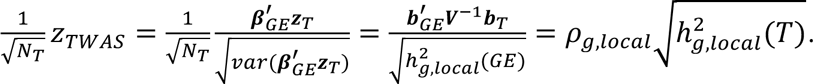

We standardize this estimate to obtain our final local genetic correlation estimate as

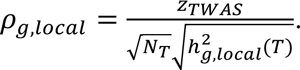

In practice we use the variance explained by the local index (i.e. smallest p-value) SNP as proxy for 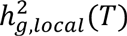.

Local components of genetic correlation characterize the shared SNP effect between complex trait and expression; however, we can interpret *ρ_g,local_* as the standardized effect of predicted expression on trait. Using this definition, we estimate the genetic correlation between two complex traits as the Pearson correlation across the vector of *ρ_g,local_* across all genes; we term this estimate as *ρ_GE_*. We test for significance assuming that 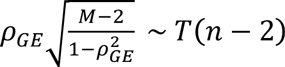 where *M* is the number of genes. This procedure is unbiased in principle provided that effects of genes within single trait are not correlated. This assumption may be violated; hence, we computed trait correlation using one gene per 1Mb locus. To determine if estimates of *ρ_GE_* were sensitive to changes in scale, we recomputed *ρ_GE_* using non-standardized estimates of genetic covariance. We found our estimates to be highly correlated (r = 0.94; p < 2.2 × 10^-16^), indicating little importance in using correlation versus covariance. We report results using standardized effects for consistency across figures and tables.

### Estimating putative casual relationships between pairs of traits

To glean insight into the underlying causal relationship between pairs of traits, we perform a bi-directional regression^14^ and estimate two different values of *ρ_GE_* by varying gene sets. Before describing the approach, we first review several causal models that explain non-zero *ρ_GE_* between two traits (see Figure 1). Models A and B depict causal relationships in which the effects of a gene set are mediated by one trait on the other. We can formally state model A (without loss of generality for B). Let *T*_1_ be defined as ***y***_*T*_1__ = ***G***_*T*_1__***β***_*T*_1__ where ***G***_*T*_1__ denotes the matrix of predicted expression at the causal genes, ***β***_*T*_1__ are the effect sizes, and *∊*_*T*_1__ is environmental noise. We define *T*_2_ as,

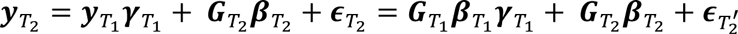

where *γ*_*T*_1__ is the causal effect of *T*_1_ on *T*_2_, *G*_*T*_1__*β*_*T*_1__ are the remaining causal genes and their effects for *T*_2_, and 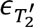 is the combined environment component. Under model A, the causal gene set for *T*_1_ will have a non-zero effect on *T*_2_ (i.e. *γ*_*T*_1__ ≠ 0); however, if *T*_1_ does not cause *T*_2_, this effect will be zero since unrelated genes have no downstream effect. Bi-directional regression provides a test to distinguish between models A and B by regressing estimated effect sizes for gene sets under model A (i.e. *β*_*T*_1__ ∼ *β*_*T*_1__*γ*_*T*_1__) and comparing to estimates under model B (i.e. *β*_*T*_2__ ∼ *β*_*T*_2__*γ*_*T*_2__). Since the causal gene sets for each trait are unknown, we use their identified susceptibility genes as proxy. We estimate *ρ_GE_* conditional on the gene set for trait *i* and denote its value as *ρ_j|i_*. This procedure is repeated by ascertaining the gene set for trait *j* to obtain *ρ_i|j_*. We perform a Welch’s t-test^44^ to determine if estimates of *ρ_i|j_* and *ρ_j|i_* are significantly different, thus providing evidence consistent with a causal direction. This approach is conceptually similar to bi-directional regression analyses of estimated SNP effects on two complex traits^15;45^. We stress that while a bi-directional approach is capable of rejecting model A in favor of model B (or vice-versa), it cannot rule out model C, in which a shared pathway (or set of pathways) drive both traits independently.

**Figure 1.**
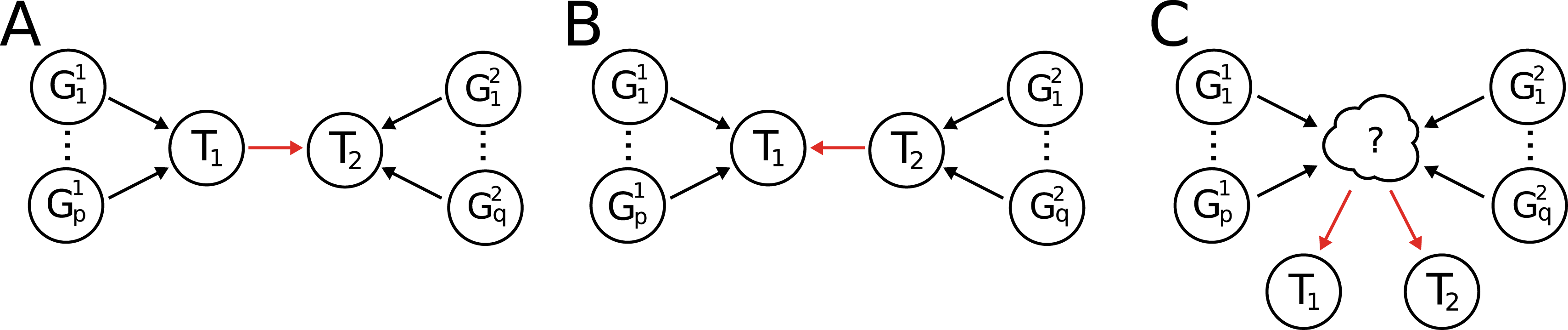
Illustration of several causal models that explain expression correlation for traits *T*_1_ and *T*_2_ given their causal gene sets. Model A) trait 1 directly influences trait 2. In this case, the effect of genes 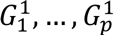 on trait 2 is mediated by trait 1 which implies 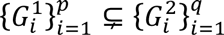. Model B) trait 2 directly influences trait 1. Similarly, the effect of genes 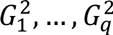 on trait 1 is mediated by trait 2 which implies 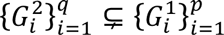. Model C) traits 1 and 2 are influenced independently through unobserved trait or traits.

## Results

### TWAS identifies 1,196 susceptibility genes for 30 complex traits and diseases

We integrated the 30 GWAS summary data with gene expression to identify 1,196 susceptibility genes (i.e. gene with at least one significant trait association) comprising 5,490 total associations (after Bonferroni correction; see Methods). Of these associations, we observed 1,789 distinct gene-trait pairs with 783 found in anthropometric traits, 423 in metabolic traits, 215 in immune-related traits, 213 in hematopoietic traits, 137 in neurological traits (i.e. schizophrenia), and 18 in social traits (see Table 1; see Supplementary Tables 3-4). For example, the 137 susceptibility genes found for schizophrenia included *SNX19* (cerebellum; p=2.2 × 10^-8^) and *NMRAL1* (muscle; p=9.7 × 10^-7^); this is consistent with a previously reported study^12^ that used different methods and expression data (see Supplementary Table 5). We did not find susceptibility genes for forearm bone mineral density (BMD), HOMA-B, and mean cell hemoglobin concentration, which is consistent with low GWAS signal for these traits (see Table 1). Indeed, the number of GWAS risk loci strongly correlated with the number of identified susceptibility genes (r=0.99; p < 2.2 × 10^-16^) which reflects the underlying polygenicity of these traits. We explored putative molecular function and pathways enriched with identified susceptibility genes using the PANTHER database^46^, but were underpowered to detect molecular function for most individual traits (see Supplementary Note).

**Table 1.**
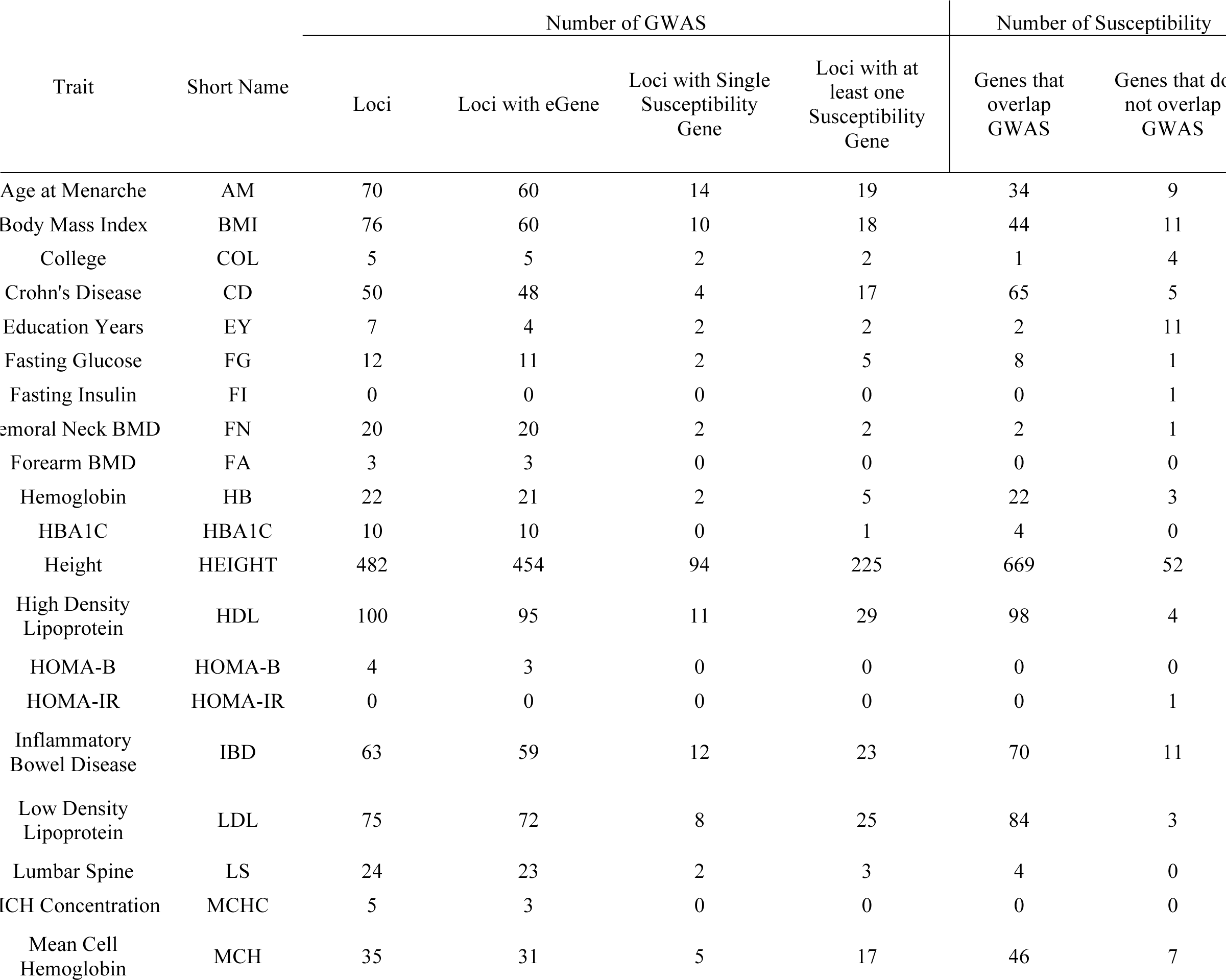
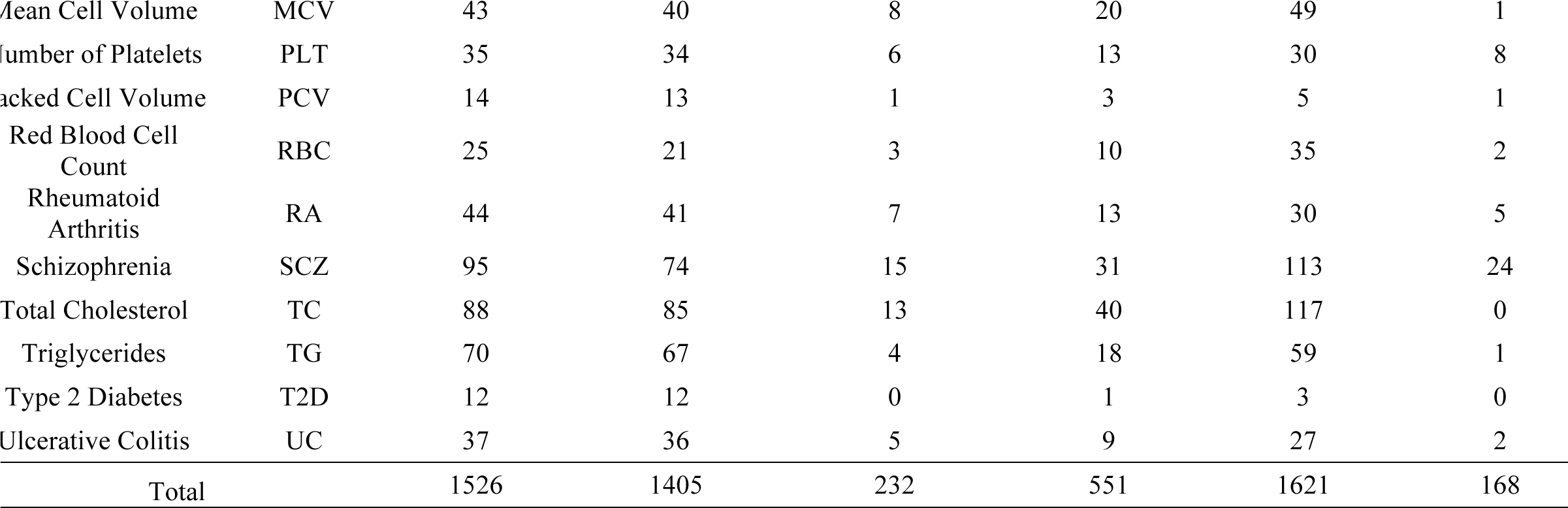
Summary of GWAS and TWAS results. The majority (92%) of GWAS risk loci overlap with at least one eGene, of which 40% contain at least one susceptibility gene. We report 168 (9%) of identified gene-trait pairs do not overlap a GWAS variant, which provide novel risk loci for follow up.

Next, we quantified the overlap of susceptibility genes and GWAS signals. Of the 1,789 identified gene-trait pairs, 168 (9%) were not proximal (more than 0.5Mb from TSS) to any genome-wide significant SNP for that respective trait thus yielding new risk loci. Conversely, of the 1,526 GWAS risk loci, 1,405 (92%) overlapped with at least one eGene (i.e. gene with heritable expression levels in at least one of the considered expression panels) and 551 (36%) overlapping at least one susceptibility gene (see Table 1). Focusing on the 1,621 associations that overlapped a genome-wide significant SNP, we observed 1,488 (83%) genes that were not nearest, suggesting that the traditional heuristic of prioritizing genes closest to GWAS SNPs is typically not supported by evidence from eQTL data (see Supplementary Figure 1). While GWAS SNPs provide the majority of the power in this approach, the flexibility of TWAS to leverage allelic heterogeneity provides a significant gain^11^. We found 219 instances across 19 traits where association signal was stronger in TWAS compared to GWAS, with an average 1.2 × increase in *χ*^2^ statistics. For example, predicted expression in *CCDC88B* (a gene involved in T-cell maturation and inflammation^47^) exhibited strong association with Crohn’s disease (p_TWAS_=6.32 × 10^-8^) whereas the index SNP (i.e. top overlapping GWAS SNP) at site rs11231774 was only suggestive (p_GWAS_=2.47 × 10^-6^). This effect was most dramatic for height, with 108 susceptibility genes having stronger signal than GWAS index SNPs. We observed a 2.6 × increase in *χ*^2^ statistics for predicted expression in *CRELD1* (p_TWAS_=1.55 × 10^-10^) compared to the index SNP rs1473183 (p_GWAS_=6.33 × 10^-5^).

Recent work^48^ applied a similar approach^12^ using summary eQTL from blood and GWAS data to identify 71 genes for 28 complex traits^48^. Of the investigated traits, 12 overlapped our study. Surprisingly, despite using independent methods and expression data we were able to validate 40 out of 51 associations (see Supplementary Table 6). Overall, we identified 564 genes for these traits in contrast to 63 genes reported in that study. This increase in power can be attributed to two reasons. First, we integrate multiple expression panels sampled from many tissues, which assays many more genes. Second, we use a method that jointly tests the entire locus, rather than index SNPs. We have shown that many identified susceptibility genes contain signals of allelic heterogeneity; therefore, using individual SNPs will decrease power.

### Genes associated to multiple traits

We investigated the degree of pleiotropic susceptibility genes (i.e. gene associated with more than one trait) in our data and found 380 (32%) identified genes associated with multiple traits (see Supplementary Figure 2). For example, the gene *IKZF3* displayed strong associations in Crohn’s disease (blood; p=1.6 × 10^-9^), HDL levels (blood; p=6.6 × 10^-15^), IBD (blood; p=7.9 × 10^-16^), rheumatoid arthritis (blood; p=6.0 × 10^-8^), and ulcerative colitis (blood; p=9.2 × 10^-10^). Indeed, *IKZF3* has been shown to influence lymphocyte development and differentiation^49; 50^. These traits are known to have a strong autoimmune component^51^; hence, association with predicted *IKZF3* expression levels is consistent with a model where cis-regulated variation in *IKZF3* product levels contributes to risk. Similarly, we observed three susceptibility genes shared between education years and height (see Figure 2): *ABCB9* (heart; p_h_eight=1.38 × 10^-15^, p_ey_=1.28 × 10^-6^), *BTN2A3P* (adipose; pheight=3.82 × 10^-12^, p_ey_=1.90 × 10^-7^), and *MPHOSPH9* (thyroid; pheight=5.84 × 10^-18^, p_ey_=1.30 × 10^-6^). This is consistent with a recent study^13^ that reported a nonzero genetic correlation between height and education years (*ρ_g_* = 0.13, p=3.82 × 10^-6^).

**Figure 2.**
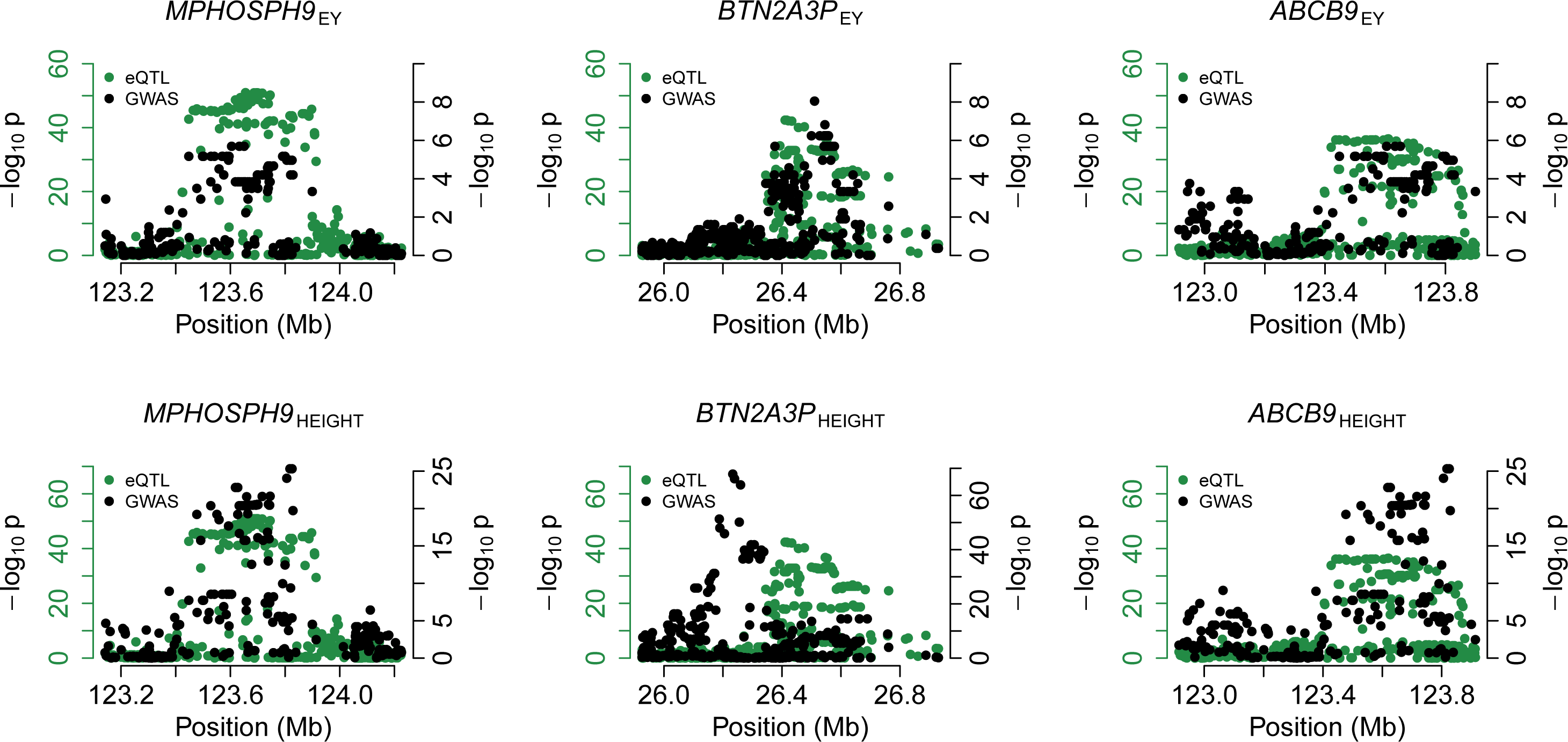
Susceptibility genes shared for education years and height. We indicate −log_10_ p-values for eQTLs in green and trait-specific GWAS in black using separate axes to simplify illustration. Their respective TWAS p-values are *ABCB9* (heart; p_h_eight=1.38 × 10^-15^, p_ey_=1.28 × 10^-6^), *BTN2A3P* (adipose; pheight=3.82 × 10^-12^, p_ey_=1.90 × 10^-7^), and *MPHOSPH9* (thyroid; pheight=5.84 × 10^-18^, p_ey_=1.30 × 10^-6^).

### Effect of cis expression on trait is consistent across tissues

Having established the importance of individual predicted gene expression levels for these traits, we next estimated the amount of trait variance explained by predicted expression using all examined genes, including those not significantly associated, using an LD-Score regression approach (see Methods). We found 108 tissue-trait pairs across 17 traits and 33 tissues where the cumulative effect of all measured genes on trait was significantly greater (p < 0.05 / 45) than the significant-only set. For example, in height we estimated 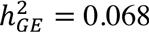 (Jack-knife SE=0.02; p=5.6 × 10^-4^; see Supplementary Table 7) using all 3,733 measured genes in YFS and 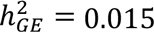 (Jack-knife SE=6.9 × 10^-3^; p=0.026) using the 169 YFS susceptibility genes (pALL>SIG=5.6 × 10^-3^). This suggests that there exist additional susceptibility genes for height, which we are underpowered to detect. However, for most trait-tissue pairs we did not observe a significant difference at our given sample sizes. Indeed, we measured a significant association between expression study sample size and number of eGenes (r=0.2; SE=0.05; p=6.4 × 10^-8^), which indicates that smaller studies lack power to find eGenes, thus underestimating the total 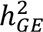.

We next asked whether any tissues are burdened with increased levels of risk for a given trait. To test this hypothesis, we examined the difference between estimated trait variance explained per gene with the average. Our results did not suggest tissue-specific enrichment at current sample sizes (see Supplementary Table 8). Given no observable difference in tissue-specific risk, we expect local estimates of genetic correlation to be highly similar across tissues. When estimating *ρ_g,local_*, we observed consistent effect size estimates in both sign and magnitude estimates across tissues (mean tissue-tissue r=0.82; see Figure 3). These results are compatible with earlier work that found cis effects on expression is largely consistent across tissues^52^. To obtain a meta estimate of local genetic correlation for gene-trait pairs with measurements in multiple tissues, we use the mean genetic correlation across all expression panels in all following analyses.

**Figure 3.**
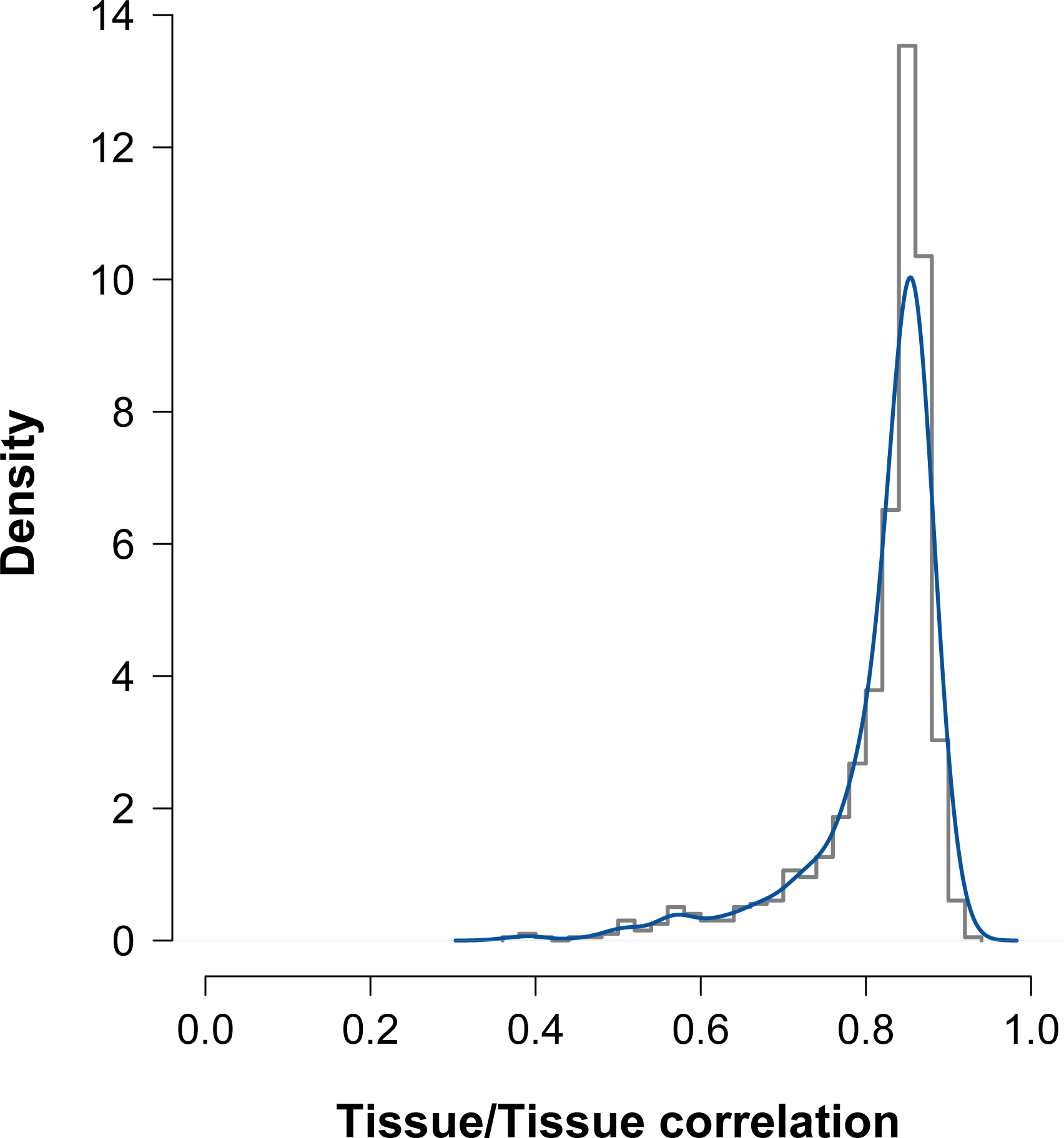
Histogram and density estimate for correlation of *ρ_g,local_* across tissues. We computed the correlation across pairs of different tissues using local estimates of genetic correlation between expression on trait. The majority of tissues exhibited high correlation over the underlying gene effects on trait with an estimated mean *r* = 0.82

### Genetic correlation between traits using predicted expression

To evaluate the shared contribution of predicted expression on pairs of traits, we computed expression correlation (*ρ_GE_*; see Methods) using nominally significant (p_TWAS_ < 0.05) genes. This approach is similar to estimating genetic correlation (*ρ_G_*) between two complex traits^13^; however, it differs in that correlation is computed through predicted components of gene expression rather than SNP effects. For 435 distinct pairs, we discovered 43 significant expression correlations, 22 of which had previously reported non-zero genetic correlations^13^ (see Figure 4; see Supplementary Table 9). For example, age of menarche and BMI had an estimated *ρ_GE_* = −0.32 (95% CI [-0.32, -0.21]; p=7.97 × 10^-8^). This negative correlation is consistent with estimates published in epidemiological studies^53^ in addition to studies probing genetic correlation across complex traits^13^. Using estimates of *ρ_GE_*, we clustered traits and observed groups forming naturally in the trait-trait matrix (see Figure 4). Interestingly, BMI clustered with insulin-related traits (HOMA-B, HOMA-IR, and fasting insulin). Our estimates were highly consistent with LD-Score regression results (see Figure 4; Supplementary Table 9). Out of 435 pairs of traits, 35 demonstrated significance for *ρ_GE_* and *ρ_g_*, whereas 8 and 27 were exclusive for *ρ_GE_* and *ρ_g_*, respectively. Given the high degree of concordance between estimates of *ρ_GE_* and *ρ_g_*, we tested if any were significantly different and found four insulin-related pairs of traits and three blood-related pairs with more extreme values for *ρ_GE_* (see Supplementary Table 9). Differences for these pairs of traits can be partially explained by overconfident standard errors in *ρ_GE_* (see Supplementary Table 10). Overall, we found *ρ_GE_* to explain the majority of variation in *ρ_g_* (r^2^ = 0.72).

**Figure 4.**
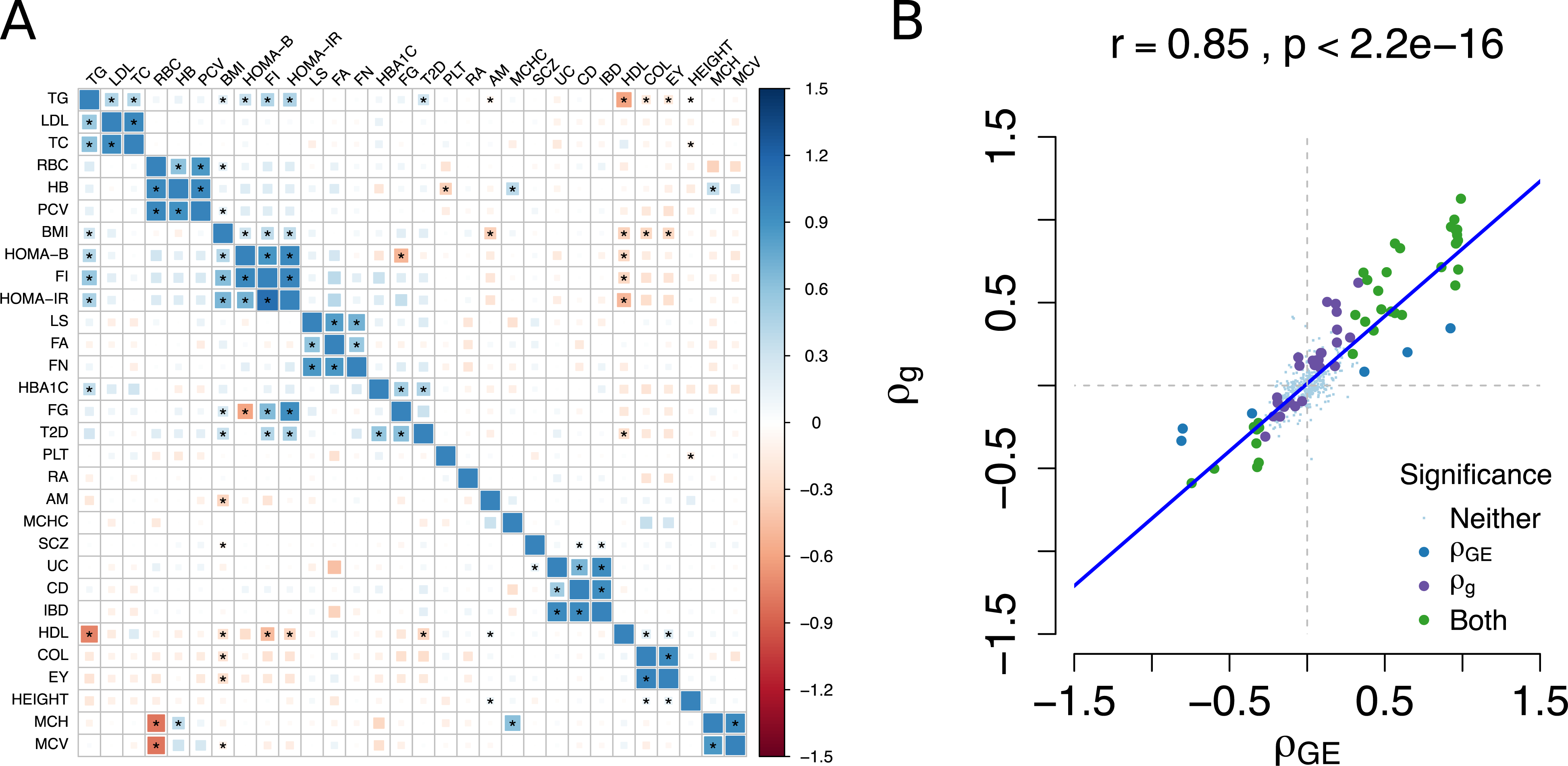
Estimates of genetic correlation *ρ_g_* obtained from LD-Score vs estimates of expression *ρ_GE_* using nominally significant TWAS results. A) Correlation matrix for 30 traits. The lower triangle contains *ρ_GE_* and the upper triangle contains *ρ_g_* estimates. Estimates of correlation that are significantly non-zero (p < 0.05 / 435) are marked with a star (*). Strength and direction of correlation is indicated by size and color. We found 43 significantly correlated traits using cis expression and 62 using genome-wide SNPs. B) Linear relationship between estimates of *ρ_GE_* and *ρ_g_*. We indicate whether individual estimates were significant in either approach by color. Non-significant trait pairs are reduced in size for visibility.

### Bi-directional regression suggests putative causal relationships

Given pairs of traits with significant estimates of *ρ_GE_*, we aimed to distinguish among possible causal explanations by performing bi-directional regression analyses (see Methods). To empirically validate our approach, we regressed HDL, LDL, and triglycerides with total cholesterol. Total cholesterol (TC) is the direct consequence of summing over triglyceride, HDL, and LDL levels, thus we expect to observe increased signal for *ρ_TC|Lipid_* compared to *ρ_Lipid|TC_*. Of these three, we found evidence for triglycerides influencing total cholesterol (p=2.34 × 10^-3^). We observed consistent, but not significant, evidence for the effect of LDL on TC (p=6.79 × 10-^2^) and HDL on TC (p=5.56× 10^-1^; see Figure 5). These results suggest that point-estimates from the bi-directional approach favor the correct model, but may not have adequate power required for significance.

**Figure 5.**
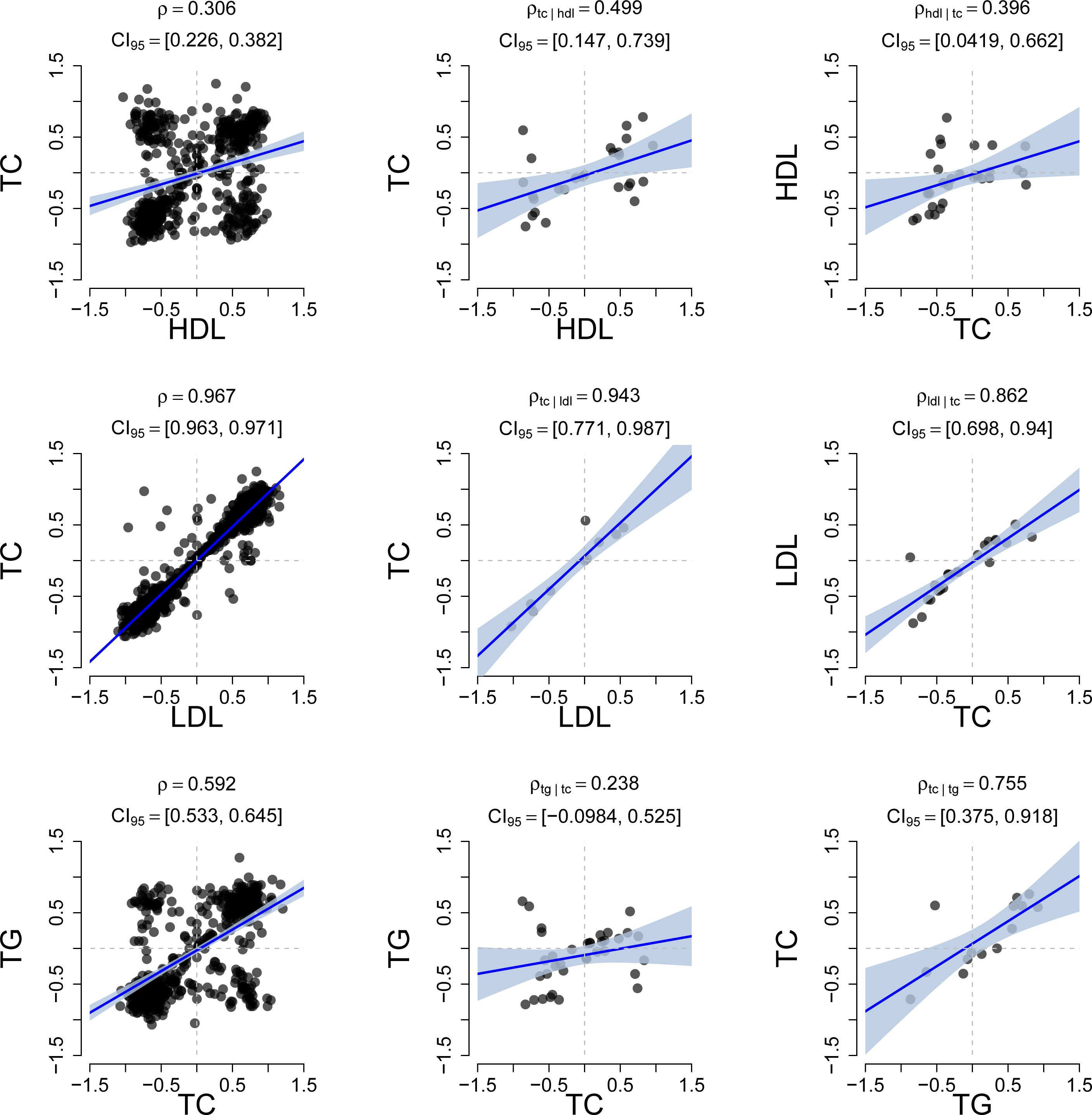
Estimates of expression correlation *ρ_GE_* for HDL, LDL, and TG with total cholesterol. Column A) Estimates of *ρ_GE_* using nominally significant genes (p < 0.05). Column B) We repeated the analysis using only susceptibility genes found in the x-axis trait but not found in the y-axis trait. Column C) Same analysis as Column B, but using the other trait’s susceptibility genes. All three analyses resulted in stronger point estimates for *ρ_TC|Lipid_* when conditioning on HDL/LDL/TG genes compared to *ρ_Lipid|TC_*; however, significance was only observed for *ρ_TC|TG_* (p=2.34 × 10^-3^).

We tested the 43 pairs of traits identified above (see Table 3) while ascertaining on susceptibility genes and observed asymmetric effects at p < 0.05 for BMI-triglycerides and LDL-triglycerides (see Figure 6). For example, in the bi-directional analysis on BMI and triglycerides, we observed a significant effect for *ρ_TC|BMI_* = 0.62 (95% CI [0.27, 0.83]; p=2.06 × 10^-3^). By contrast, the reverse analysis estimate overlapped with zero at *ρ_BMI|TG_* = −0.04 (95% CI [-0.49, 0.42]; p=0.86). Individual estimates for *ρ_TG|BMI_* and *ρ_BMI|TG_* were significantly different (p=0.01; Welch’s t-test), which is consistent with a model where BMI directly influences triglyceride levels. In practice, we used susceptibility genes found through TWAS (p ∼ 1 × 10^-6^), but this may be too strict an inclusion threshold for genes which we lack power to detect. We report analyses using weaker thresholds and observe similar results (see Supplementary Tables 11, 12). Our result reinforces previous estimates of putative causal effect where BMI influences triglyceride levels^15; 54^.

**Figure 6.**
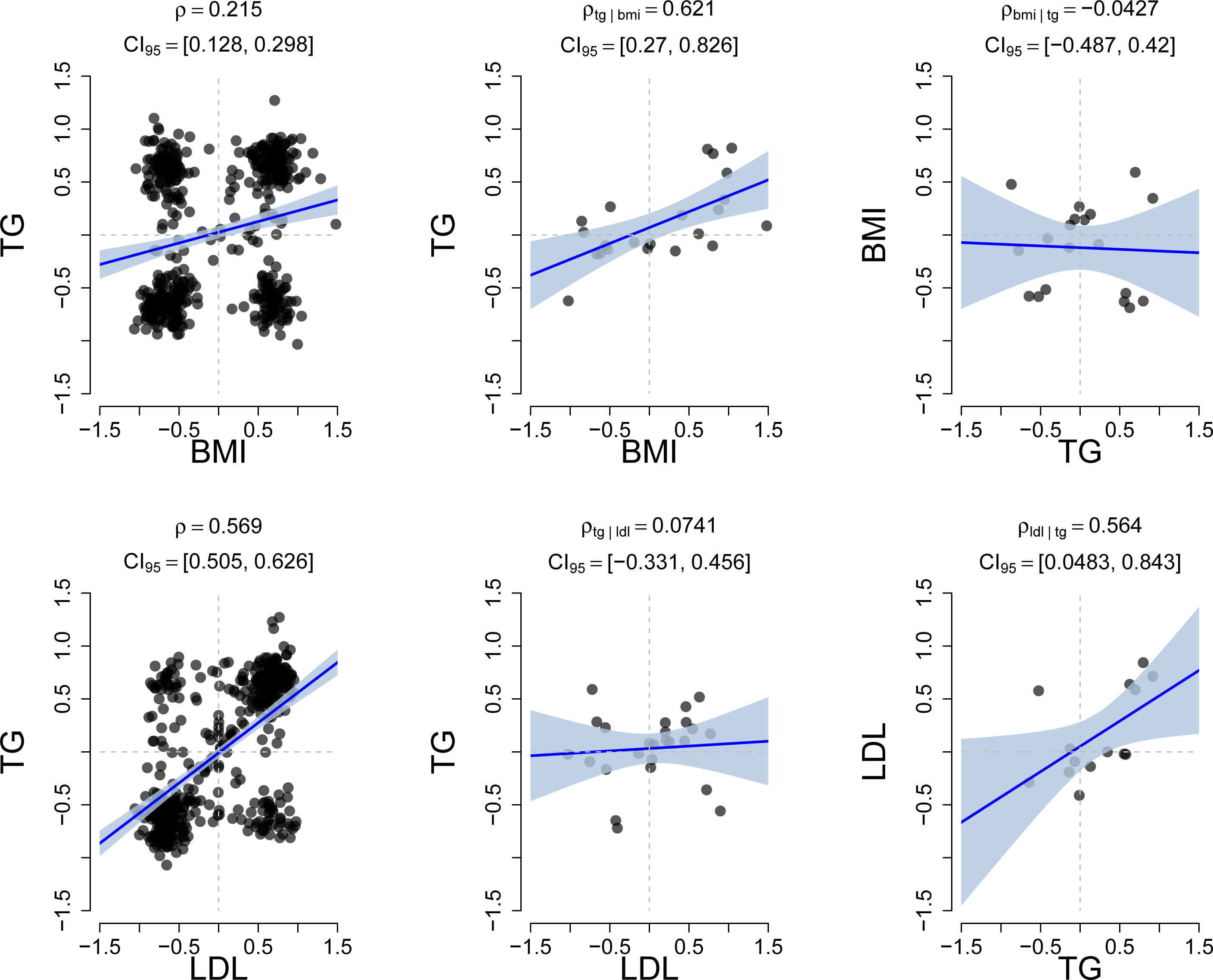
Estimates of expression correlation *ρ_GE_* for triglycerides with BMI and triglycerides with LDL. We present results for pairs of traits that displayed a significant difference (p < 0.05; Welch’s t-test) in their conditional estimates. These results are consistent with a causal model where BMI influences TG and TG influences LDL.

**Table 2.**
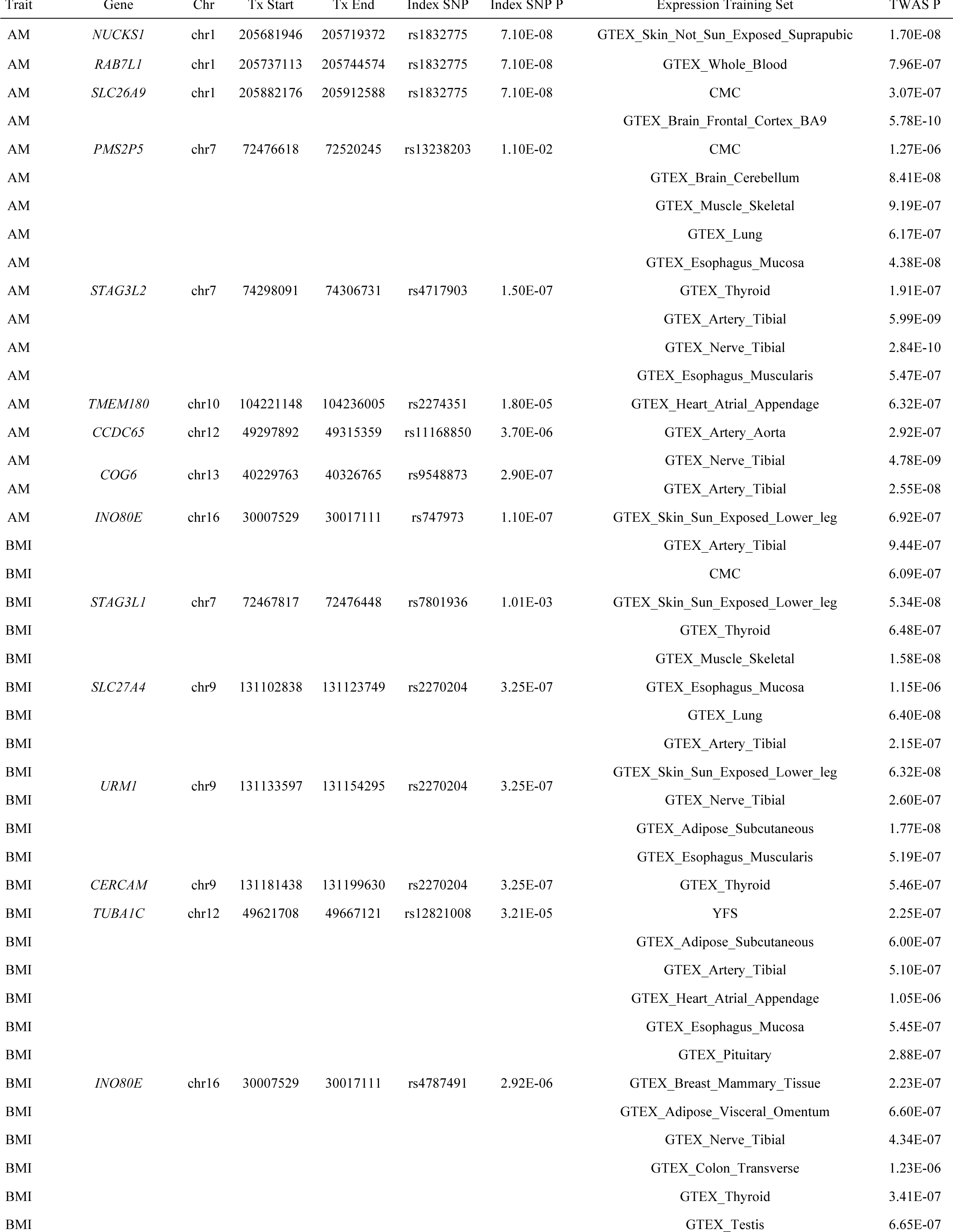
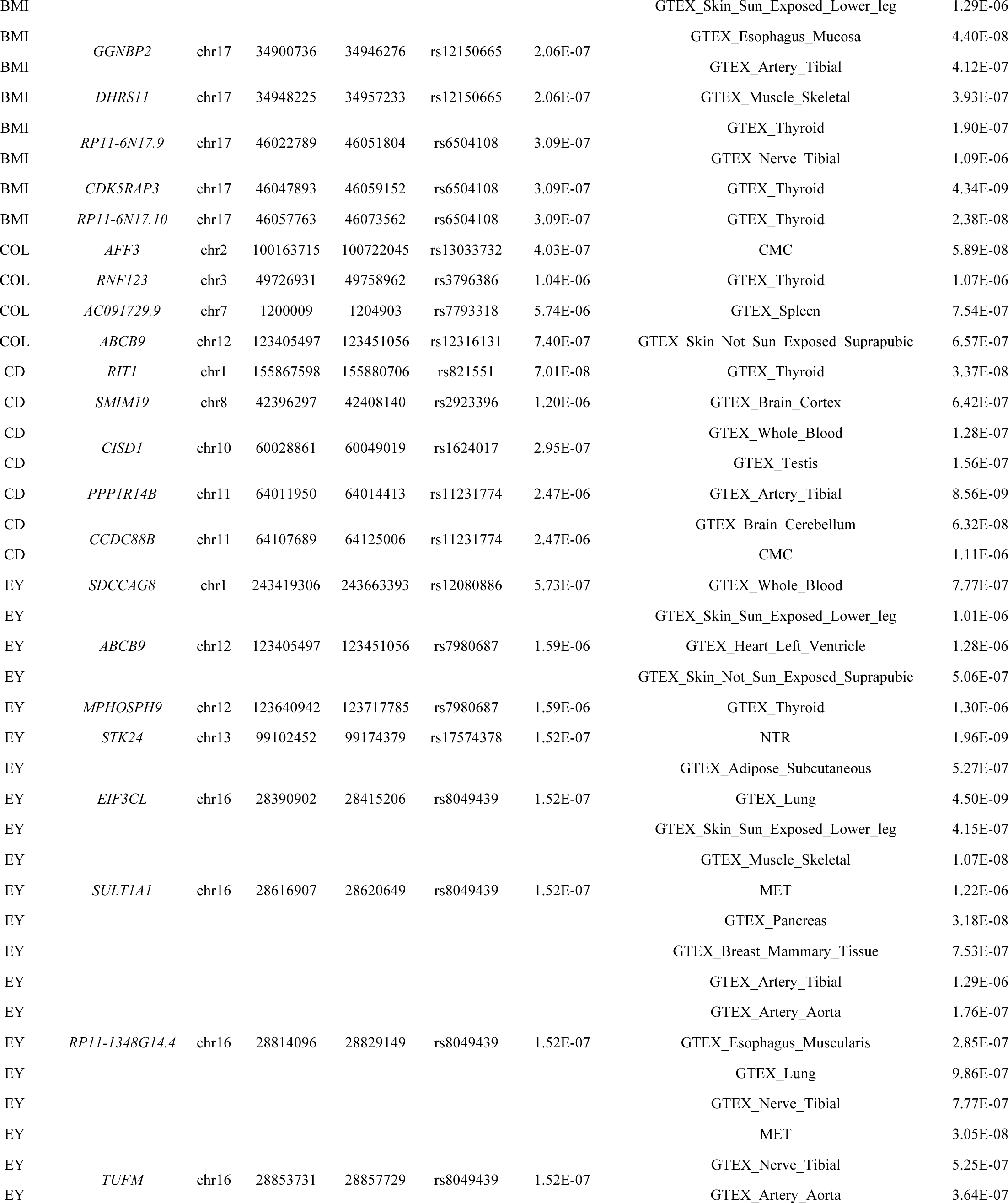
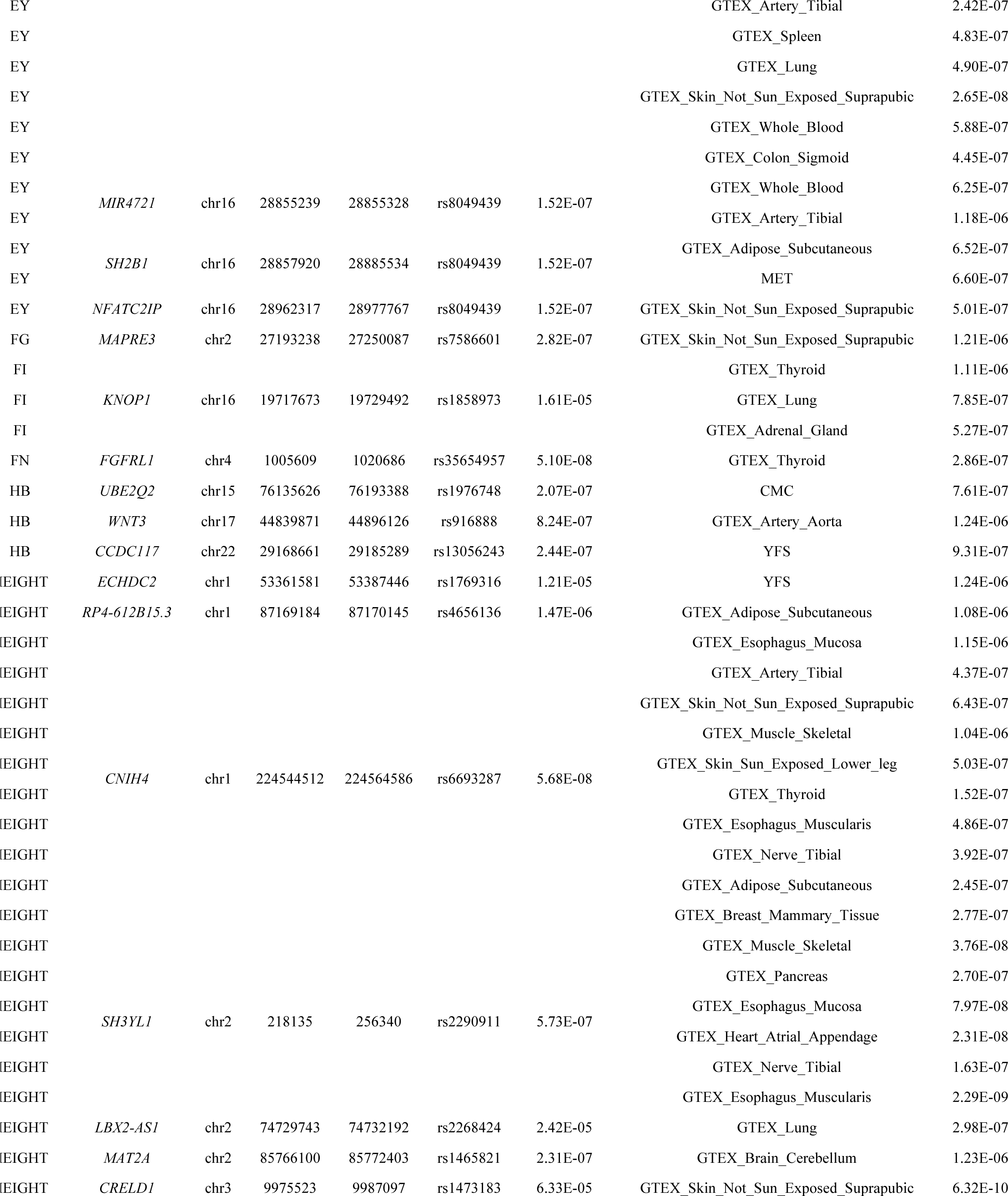
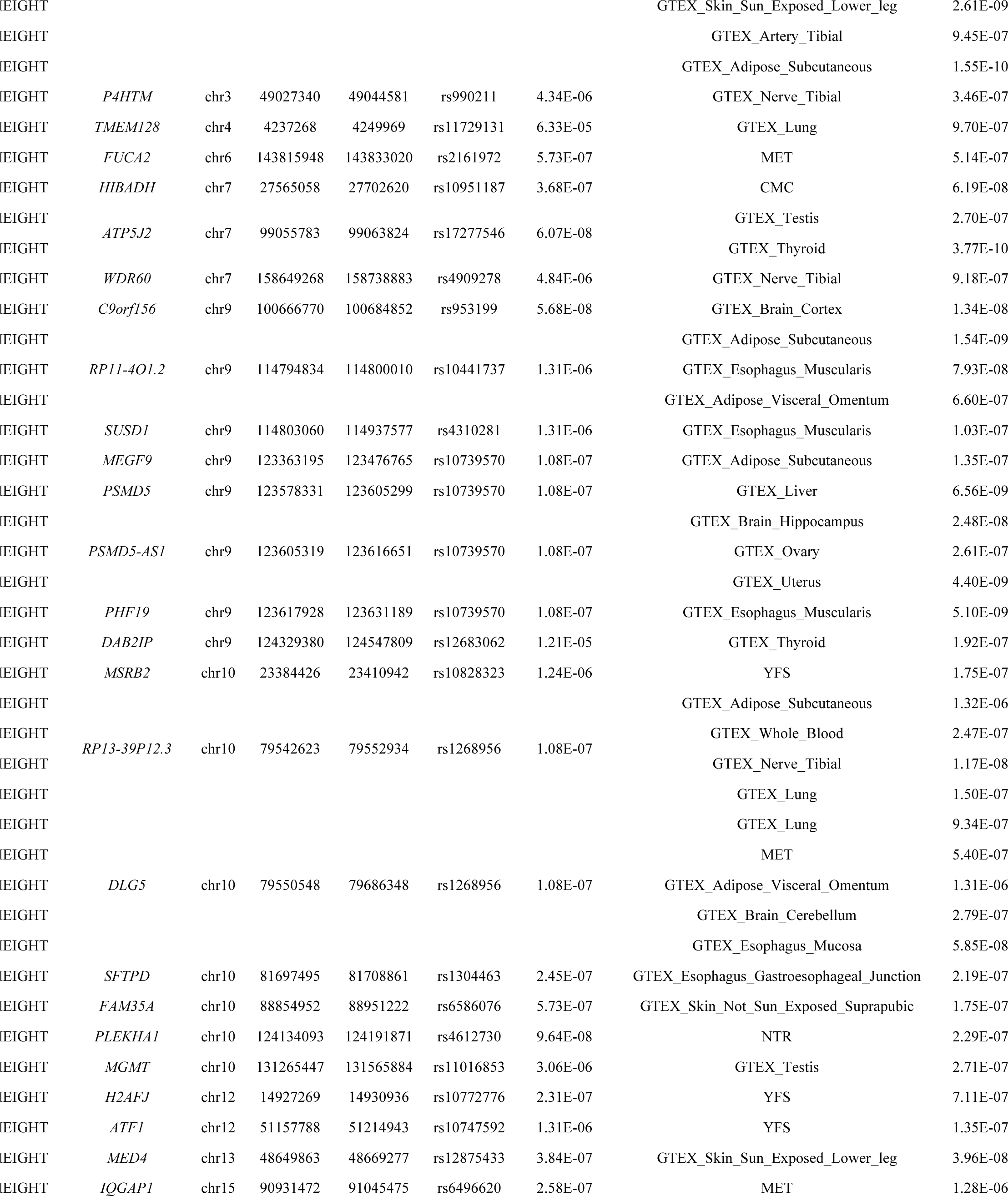
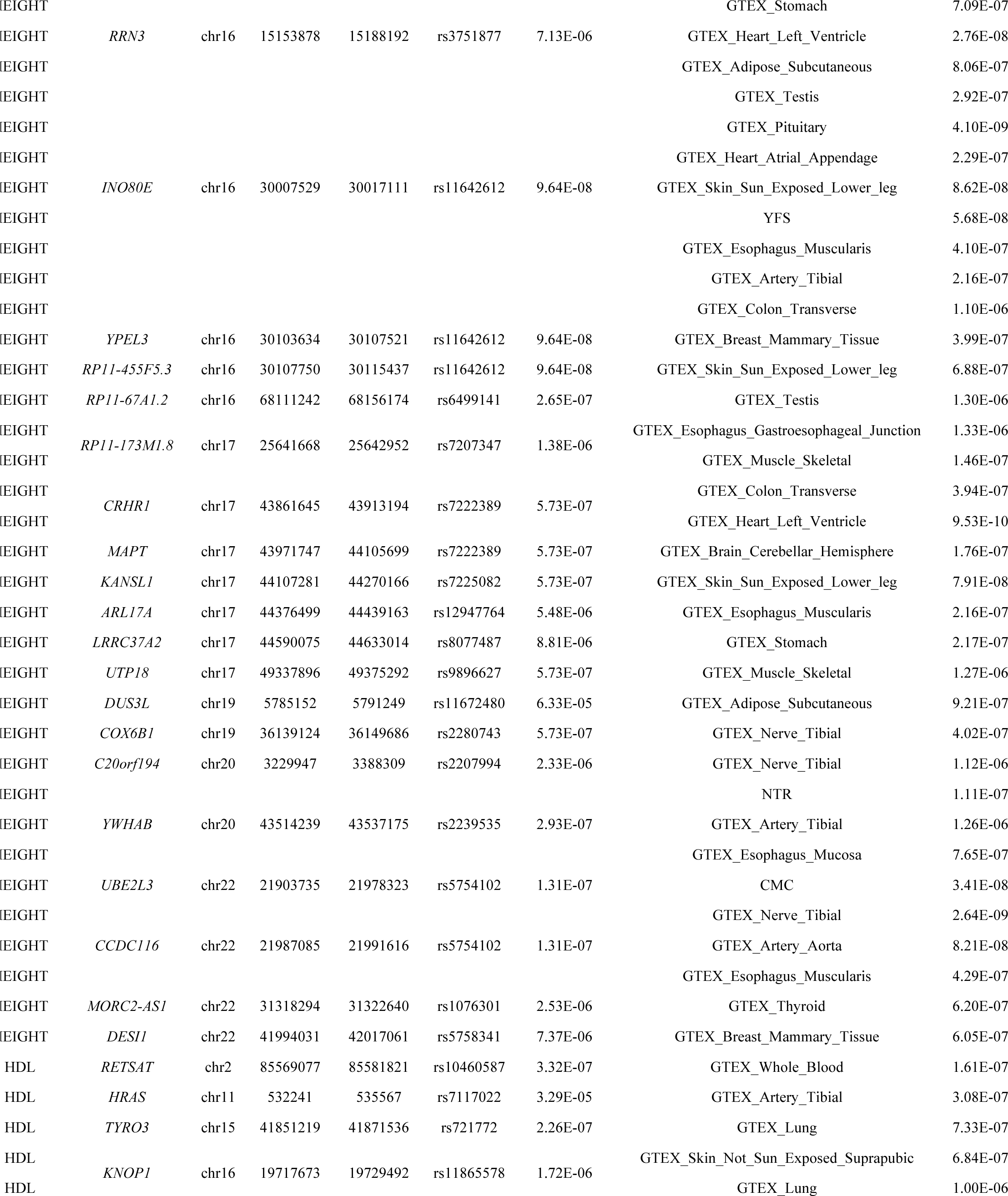
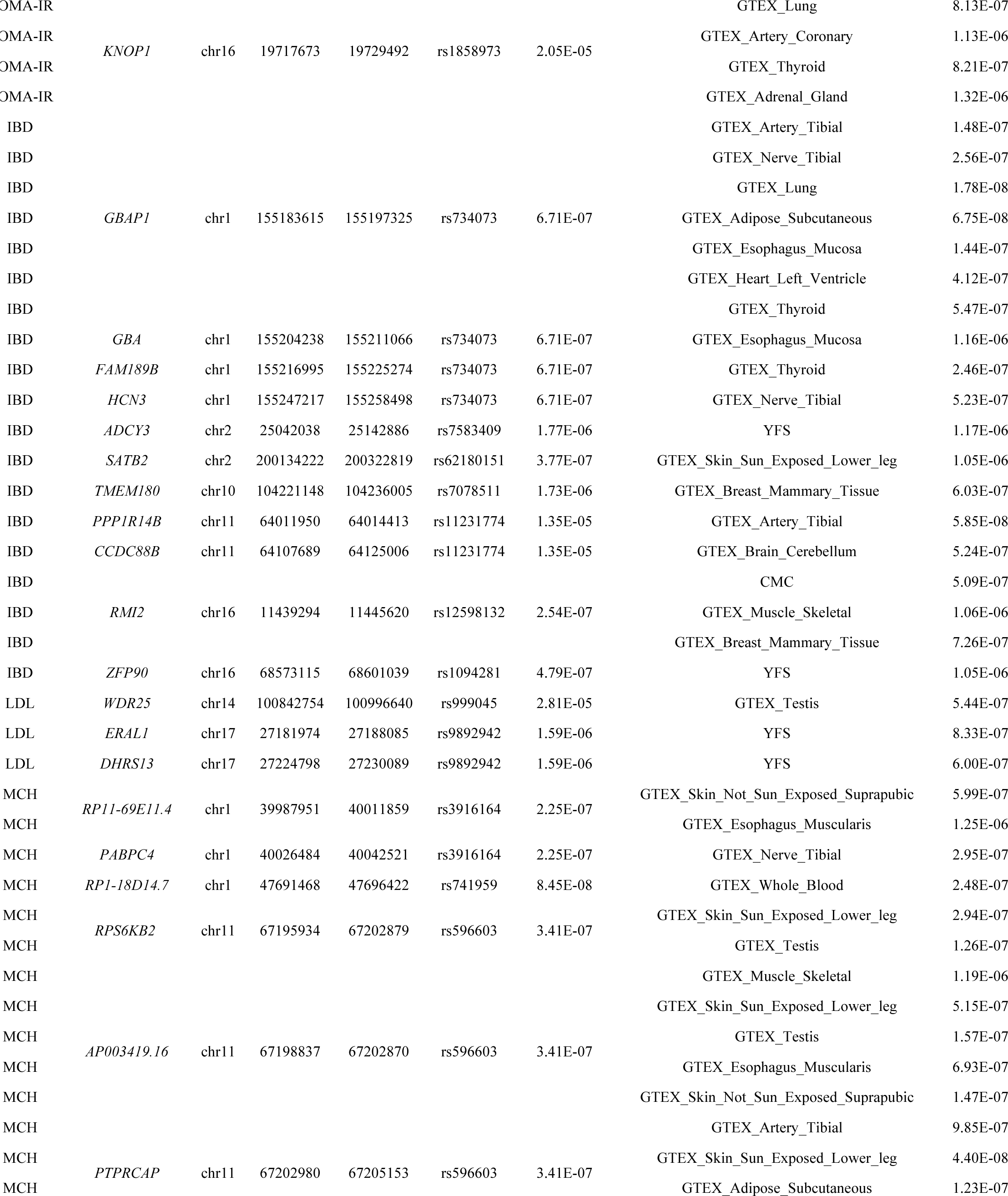
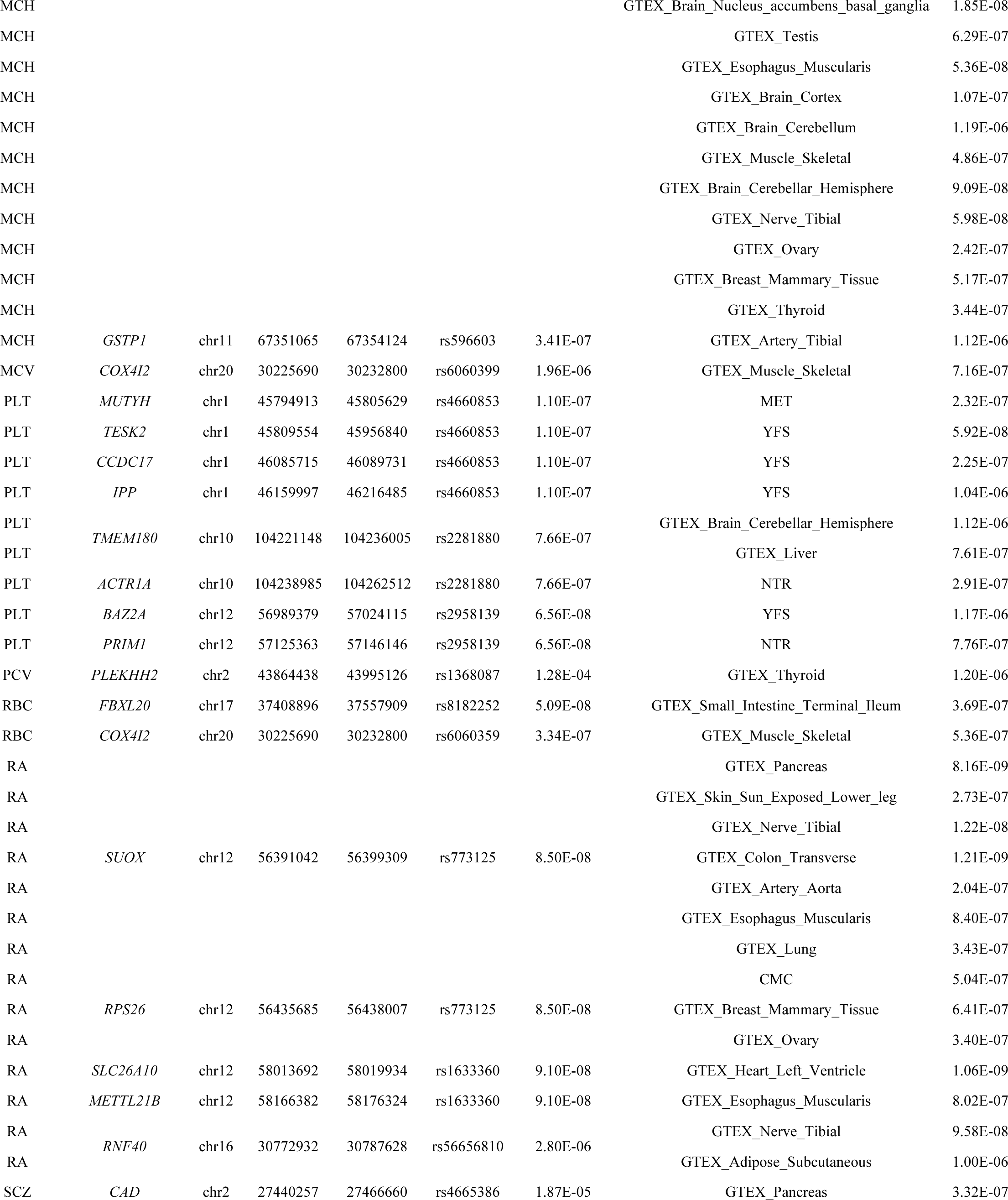
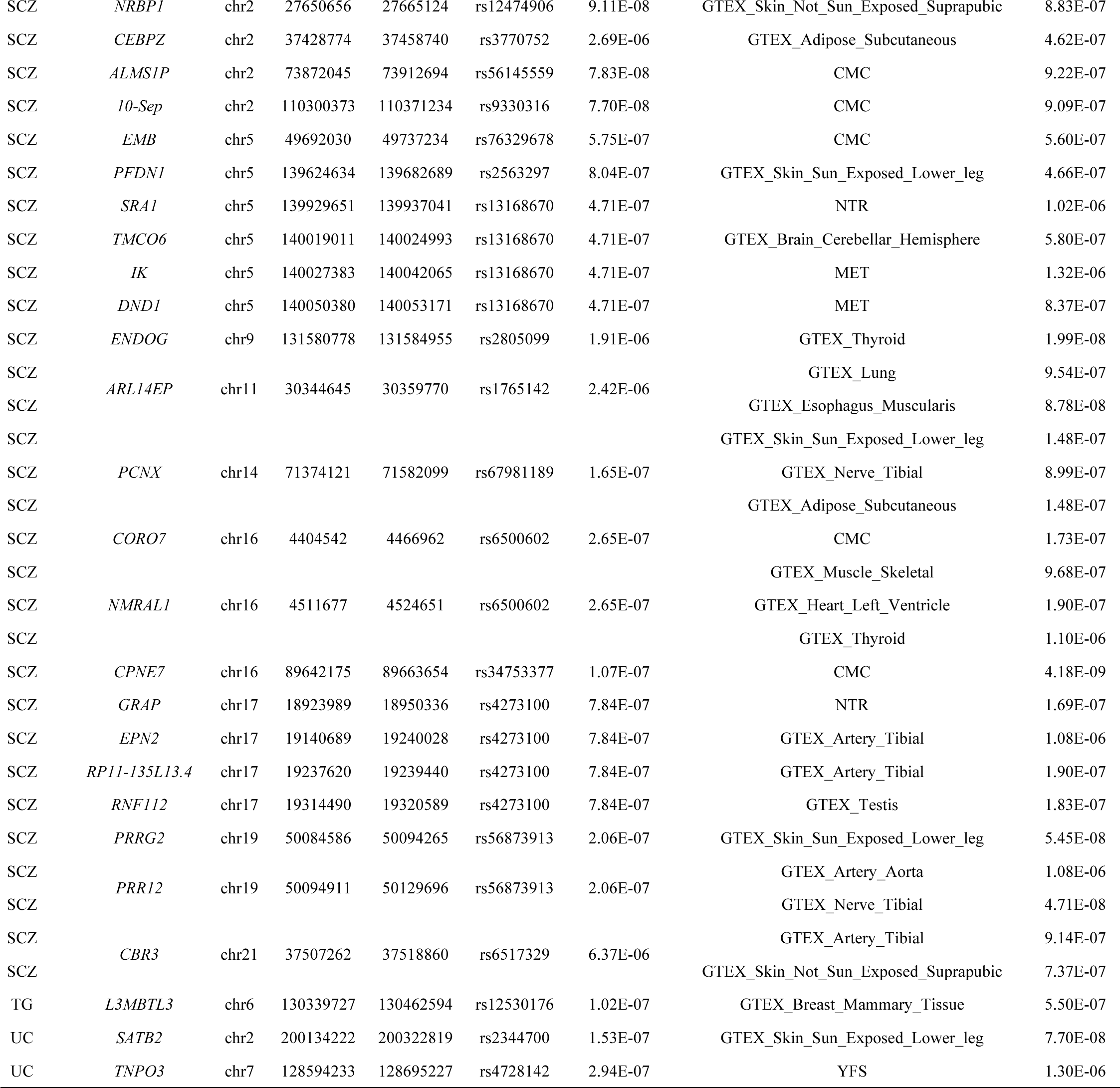
Novel risk loci. Identified susceptibility genes that do not overlap a genome-wide significant SNP (p < 5 × 10^-8^) within 0.5Mb for the tested trait.

**Table 3.**
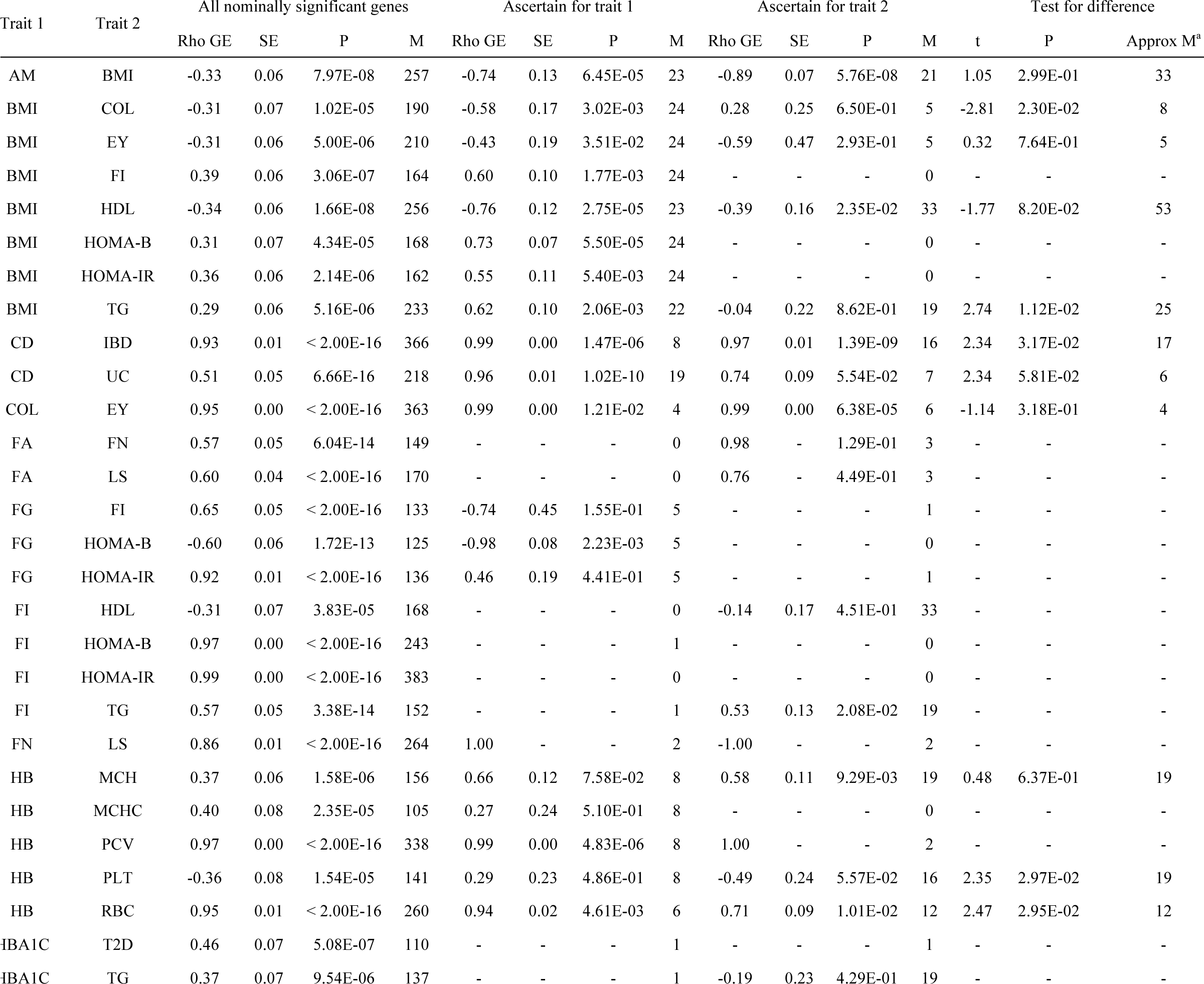
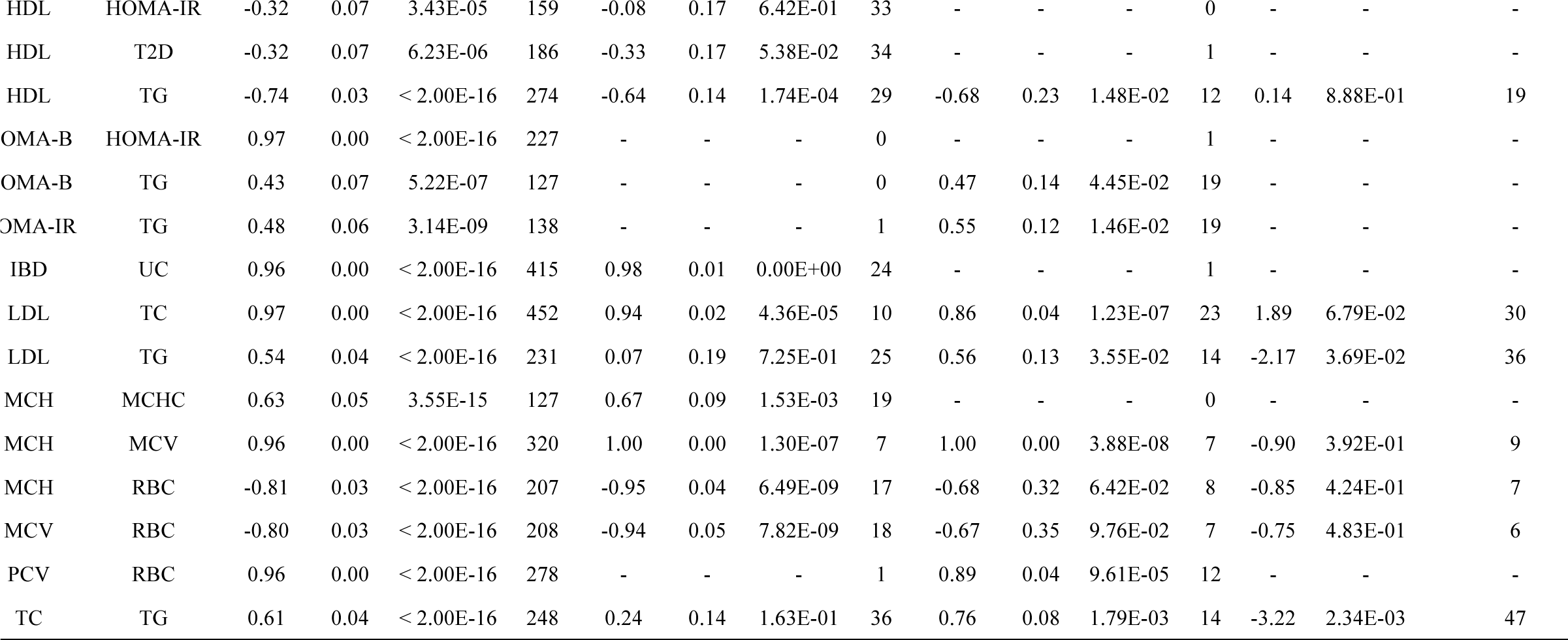
Significant estimates of *ρ_GF_* for 43 pairs of traits. We performed bi-directional regression and obtain conditional estimates of *ρ_GF_*, which provides evidence for a putative causal direction. We observed three pairs of traits with a significant difference between their directional estimates; namely, BMI influencing TG, TG influencing LDL, and TG influencing TC. We mark entries with *M* < 3 with “-“. a determined by Welch–Satterthwaite equation

## Discussion

In this work we used GWAS summary statistics from 30 complex traits and diseases jointly with expression data sampled across 45 expression panels to identify susceptibility genes for complex traits. We identified 1,196 susceptibility genes for 27 of the 30 complex traits. We use estimates of local genetic correlation between gene expression and trait to compute *ρ_GE_*, which quantifies the shared effect of predicted expression levels between two complex traits. Using this definition, we found 43 pairs of traits to be significantly correlated, of which 8 were novel. To provide evidence of possible causal direction, we adapted a recently proposed causality test^15^ to operate at the gene level. Our results support triglycerides (TG) influencing LDL, and BMI influencing triglycerides. As more GWAS and eQTL summary results become publicly available, we expect additional studies to integrate cross-trait information to make inferences about mechanistic bases for complex trait.

Assuming gene expression mediates the effect of genetics on complex trait, testing for association between the predicted component of expression and trait is equivalent with a two-sample Mendelian randomization test for a causal effect of expression on trait^55; 56^. This test for causality is valid provided SNPs do not exhibit pleiotropic effects; therefore, the TWAS associations are not proof of causal relationships between expression and complex trait. This set of assumptions extends to our bi-directional approach to infer causal direction. A bi-directional regression is capable of distinguishing between direction of effect, but cannot rule out pleiotropy.

We conclude with several caveats. First, we note that using estimates of genetic correlation to find susceptibility genes may still be biased due to confounding. The expression weights used for TWAS may tag variants that are causal through other genes or non-genic mechanisms. In principle, this can be partially remedied by jointly testing multiple genes with trait; however, a correctly specified model would require covariance estimates between observed, not predicted, expression levels—which is not available in summary data. In this work we combined estimates across tissues by taking the mean effect to compute the genetic correlation between trait and expression. This approach is unbiased, but may be inefficient. Recent work^57^ describes a random-effects model to combine estimates across tissues to increase power. Finally, our method to estimate correlation between traits using the genetically predicted component of gene expression makes several simplifying assumptions. We remedied the non-independence of genes by sampling single genes within a 1Mb region, an approach which has been used previously^45^. However, a more powerful approach may take correlations across genes into account.

## Acknowledgements

We would like to thank Valerie Arboleda, Robert Brown, Kathy Burch, and Malika Kumar for helpful discussions and feedback. We also thank Dr. Nicole Soranzo for sharing summary data for the platelet traits.

CMC: Data were generated as part of the CommonMind Consortium supported by funding from Takeda Pharmaceuticals Company Limited, F. Hoffman-La Roche Ltd and NIH grants R01MH085542, R01MH093725, P50MH066392, P50MH080405, R01MH097276, RO1-MH-075916, P50M096891, P50MH084053S1, R37MH057881 and R37MH057881S1, HHSN271201300031C, AG02219, AG05138 and MH06692. Brain tissue for the study was obtained from the following brain bank collections: the Mount Sinai NIH Brain and Tissue Repository, the University of Pennsylvania Alzheimer’s Disease Core Center, the University of Pittsburgh NeuroBioBank and Brain and Tissue Repositories and the NIMH Human Brain Collection Core. CMC Leadership: Pamela Sklar, Joseph Buxbaum (Icahn School of Medicine at Mount Sinai), Bernie Devlin, David Lewis (University of Pittsburgh), Raquel Gur, Chang-Gyu Hahn (University of Pennsylvania), Keisuke Hirai, Hiroyoshi Toyoshiba (Takeda Pharmaceuticals Company Limited), Enrico Domenici, Laurent Essioux (F. Hoffman-La Roche Ltd), Lara Mangravite, Mette Peters (Sage Bionetworks), Thomas Lehner, Barbara Lipska (NIMH)

## Web Resources

TWAS: http://bogdan.bioinformatics.ucla.edu/software/twas/

CMC: https://www.synapse.org/cmc/

GTEx: http://www.gtexportal.org/home/

GCTA: http://cnsgenomics.com/software/gcta/

